# GABAergic deficits and schizophrenia-like behaviors in a mouse model carrying patient-derived neuroligin-2 R215H mutation

**DOI:** 10.1101/225524

**Authors:** Dong-Yun Jiang, Zheng Wu, Yi Hu, Siu-Pok Yee, Gong Chen

**Affiliations:** Department of Biology, Huck Institutes of Life Sciences, Pennsylvania State University, University Park, PA 16802; Department of Cell Biology, University of Connecticut Health center, Farmington, CT, 06030

**Keywords:** Schizophrenia, GABA, neuroligin-2, mouse model, mutation

## Abstract

Schizophrenia (SCZ) is a severe mental disorder characterized by delusion, hallucination, and cognitive deficits. We have previously identified from schizophrenia patients a loss-of-function mutation Arg^215^ → His^215^ (R215H) of *neuroligin 2 (NLGN2)* gene, which encodes a cell adhesion molecule critical for GABAergic synapse formation and function. Here, we generated a novel transgenic mouse line with neuroligin-2 (NL2) R215H mutation, which showed a significant loss of NL2 protein, reduced GABAergic transmission, and impaired hippocampal activation. Importantly, R215H KI mice displayed anxiety-like behaviors, impaired pre-pulse inhibition (PPI), cognition deficits and abnormal stress responses, recapitulating several key aspects of schizophrenia-like behavior. Our results demonstrate a significant impact of a single point mutation NL2 R215H on brain functions, providing a novel animal model for the study of schizophrenia and neuropsychiatric disorders.

---

Schizophrenia (SCZ) is a chronic neuropsychiatric disorder caused by both genetic and environmental factors. It is featured by long-standing delusion and hallucination (psychosis), and cognitive deficits (Freedman, 2003; Insel, 2010; Lewis and Lieberman, 2000). SCZ is a highly heritable disorder (Sullivan et al., 2003) with a complex genetic basis. Recent genomic studies identified a number of genetic variants associated with SCZ, including a group of variants resided in the genes encoding synaptic adhesion molecules that promoting synaptic function and development such as *IGSF9B,* and *NLGN4X* (Schizophrenia Working Group of the Psychiatric Genomics, 2014).

Neuroligins (NLGNs) are a family of synaptic adhesion molecules highly expressed in the brain and are ligands for another group of cell adhesion molecules neurexins (NRXNs) (Ichtchenko et al., 1995). There are five neuroligin genes (neuroligin-1, -2, -3, -4, and -5) in humans and four in mice (neuroligin 1-4). Neuroligin-1, -2, and -3 are close homologs between human and mice. Neuroligin-1 and neuroligin-2 differentially locate to excitatory and inhibitory synapses and are critical for the excitatory and inhibitory synapse formation and function, respectively (Chubykin et al., 2007; Dong et al., 2007; Levinson et al., 2005; Nam and Chen, 2005; Scheiffele et al., 2000; Song et al., 1999; Varoqueaux et al., 2004). Neuroligin-3 locates at both type of synapses and contributes to both neurotransmission (Budreck and Scheiffele, 2007; Etherton et al., 2011). In recent years, several genetic variants of neuroligin-3 and neuroligin-4 have been identified in autism patients (Chih et al., 2004; Jamain et al., 2003). Mutations in proteins interacting with neuroligins such as Neurexin1, SHANK and MDGA have also been associated with autism and schizophrenia patients (Bucan et al., 2009; Durand et al., 2007; Kim et al., 2008; Kirov et al., 2008). Genetic mouse models based on these findings recapitulate some aspects of patient symptoms as well (Baudouin et al., 2012; Connor et al., 2016; Etherton et al., 2011; Etherton et al., 2009; Jamain et al., 2008; Peça et al., 2011; Rothwell et al., 2014; Südhof, 2008; Tabuchi et al., 2007; Zhou et al., 2016).

We have previously reported several novel mutations of *NLGN2* from schizophrenia patients (Sun et al., 2011). Among the NL2 mutants, we found that the R215H mutant protein was retained in the endoplasmic reticulum (ER) and could not be transported to the cell membrane, resulting in a failure to interact with presynaptic neurexin and a loss of function in GABAergic synapse assembly (Sun et al., 2011). Based on these studies, we have now generated a transgenic mouse line carrying the same NL2 R215H mutation to test its functional consequence *in vivo.* We demonstrate that the R215H knock-in (KI) mice show severe GABAergic deficits and display anxiety-like behavior, impaired pre-pulse inhibition, cognitive deficits, and abnormal stress responses. These deficits are more severe than reported NL2 KO mice. Our results suggest that a single-point mutation R215H of NL2 can result in significant GABAergic deficits and contribute to SCZ-like behaviors. This newly generated NL2 R215H KI mouse may provide a useful animal model for the studies of neuropsychiatric disorders including SCZ.

## Results

### Generation of neuroligin-2 R215H mutant mice

Following our original discovery of a loss-of-function mutation R215H of NL2 in SCZ patients (Sun et al., 2011), we generated the NL2 R215H mutant mice by introducing the same R215H mutation into the exon 4 of *Nlgn2* gene in the mouse genome via homologous recombination (Figure 1a). NL2 R215H heterozygotes were mated to obtain wild type (WT), heterozygotes (referred here as Het mice), and homozygotes (referred here as KI mice) (Figure S1a). Sequencing analysis confirmed the R215H mutation in the NL2 KI mice (Figure S1b). Mice carrying R215H mutation were born at a normal Mendelian rate (Male mice: WT = 26.5%, Het = 52.9%, KI = 20.6%; Female mice: WT = 24.1%, Het = 52.8%, KI = 23.1%). Both R215H Het and KI mice were viable and fertile and did not exhibit premature mortality. The weight of R215H Het and KI mice was not significantly different from WT mice (Figure S2). The mouse colony was maintained on a hybrid genetic background to avoid the artificial phenotype contributed by other homozygous genetic variants in a homozygous inbred background.

**Figure 1.**
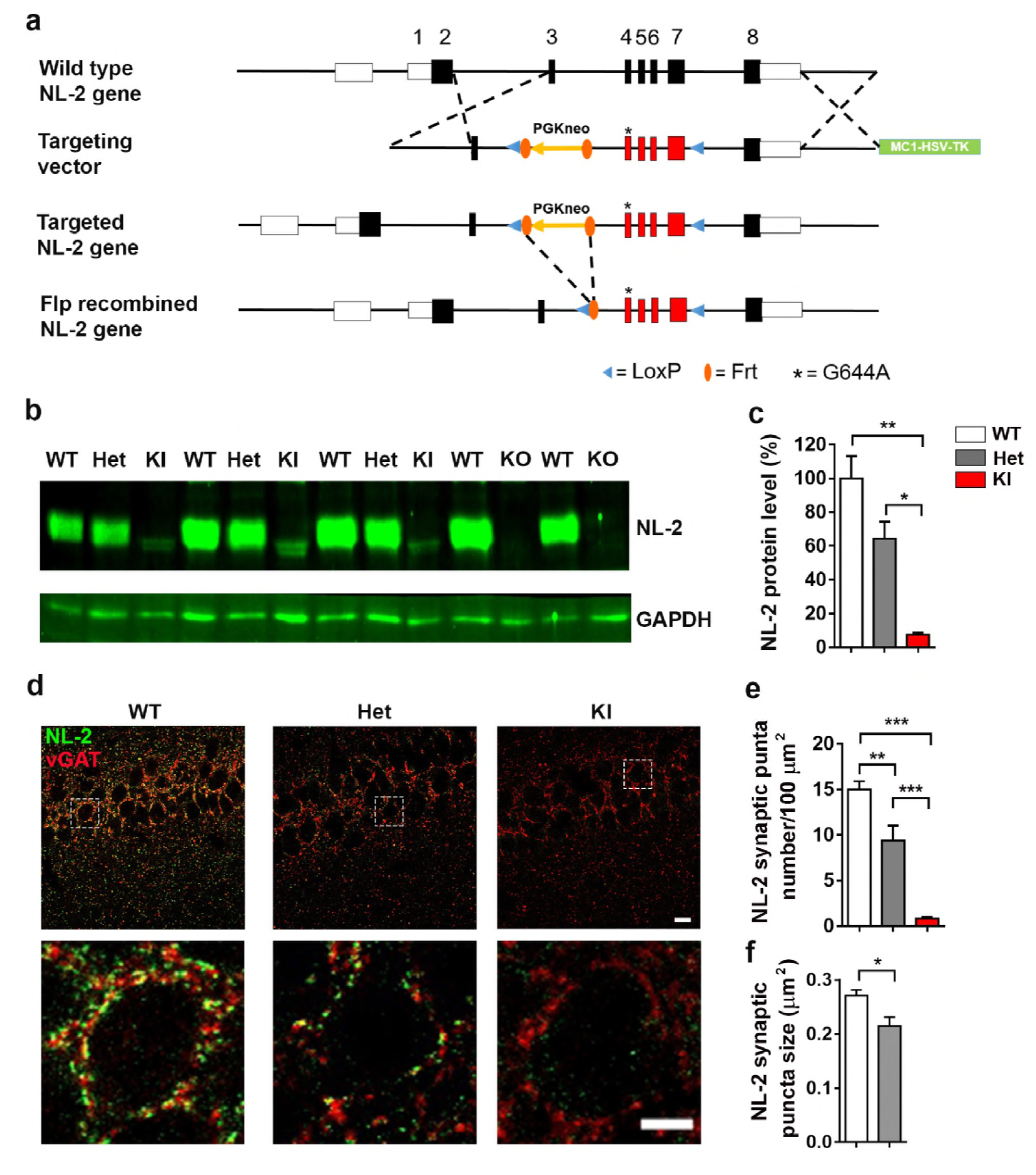
Generation and characterization of NL2 R215H mice. **(a)** A simplified diagram of NL2 R215H homologous recombination strategy. **(b)** Representative Western blot of NL2 protein expression in total brain homogenates of WT, NL2 R215H Het, and NL2 R215H KI mice. GAPDH was used as internal control. **(c)** Quantification of NL2 expression level. 3 repeats of WT, Het, and KI littermates were used for quantification. One-way ANOVA with post-hoc Tukey multi-comparison test was used for statistical analysis. **(d)** Representative images of NL2 postsynaptic puncta in WT, NL2 R215H Het and NL2 R215H KI mice. Images were taken at hippocampal CA1 region. Upper row scale bar = 10 μm. Bottom row scale bar = 5 μm. **(e-f)** Quantification of NL2 puncta number and size. 9 brain slices from 3 mice for each genotype were used for analysis. One-way ANOVA with post-hoc Tukey multicomparison test was used for statistical analysis in **(e)**. Student’s *t*-test was used for statistical analysis in **(f)**. Data were shown as Mean ± SEM, *P < 0.05, **P < 0.01, ***P < 0.001.

### Reduction of neuroligin-2 protein level in NL2 R215H KI mice

After obtaining the NL2 R215H Het and KI mice, we first analyzed the NL2 protein expression level in the brain. We found that NL2 expression level reduced 90% in NL2 R215H homozygotes, and 40% in heterozygotes (Figures 1b and c), which was in contrast to a complete absence of NL2 in the NL2 KO mice (Figure 1b). Notably, the residual NL2 R215H protein band only showed a very weak lower band compared to the thick WT NL2 protein bands, suggesting that the residual NL2 R215H proteins were likely immature NL2 without glycosylation (Sun et al., 2011; Zhang et al., 2009).

To investigate the localization of NL2 R215H proteins inside the brain, we performed immunohistochemistry with NL2-specific antibodies and found a significant reduction of NL2 puncta in R215H Het mice and almost absence of NL2 puncta in homozygous R215H KI mice (Figure 1d-f). In WT mouse brains, NL2 formed numerous postsynaptic puncta on cell soma and dendrites opposing presynaptic vGAT puncta (Figure 1d-f, puncta density 15.0 ± 0.9 per 100 μm^2^, puncta size = 0.27 ± 0.01 μm^2^).

The number and size of NL2 puncta were significantly reduced in the NL2 R215H Het mouse brains (Figure 1d-f, puncta density, 9.4 ± 1.6 per 100 μm^2^, p < 0.01, One way ANOVA followed with Tukey post hoc test; puncta size, 0.21 ± 0.02 μm^2^, p = 0.02). Interestingly, in the homozygous NL2 R215H KI mouse brains, only faint NL2 signal was observed inside cell soma (Figure 1d, right columns) and not colocalized with vGAT, indicating that the NL2 R215H proteins could not be transported to the cell membrane, consistent with our previous observation in cell cultures (Sun et al., 2011). To get a clear understanding of the physiological role of NL2 R215H mutation in *vivo*, we focused our studies on the homozygous NL2 R215H KI mice in this study.

### Reduced GABAergic synapse density in NL2 R215H KI mice

NL2 has been reported to form complex with gephyrin and collybistin at postsynaptic sites to recruit GABA_A_ receptors (Poulopoulos et al., 2009). Consistent with a substantial reduction of NL2 puncta in the KI mice, we detected a remarkable decrease of postsynaptic GABA_A_ receptor γ2 subunit and the scaffold protein gephyrin around cell soma in hippocampal regions (Figure 2a). Quantitative analysis revealed that both the puncta number and size of postsynaptic γ2 subunit and gephyrin decreased significantly in homozygous R215H KI mice (Figure 2b-e), consistent with previous findings in NL2 KO mice (Babaev et al., 2016; Gibson et al., 2009; Jedlicka et al., 2010; Poulopoulos et al., 2009). In addition to postsynaptic changes, we also observed a significant reduction of presynaptic PV and vGAT puncta (vesicular GABA transporter) in the hippocampal region of KI mice (Figure 3, Figure S3). The number of PV neurons was not changed in the KI mice (Figure 3b-d). However, both PV and vGAT puncta number and size were significantly reduced in the dentate granule cells (Figure 3e-i), as well as in the CA1/CA3 pyramidal cells in KI mice (Figure S3a-j). These results suggest that NL2 R215H mutation impaired both pre- and post-synaptic GABAergic components.

**Figure 2.**
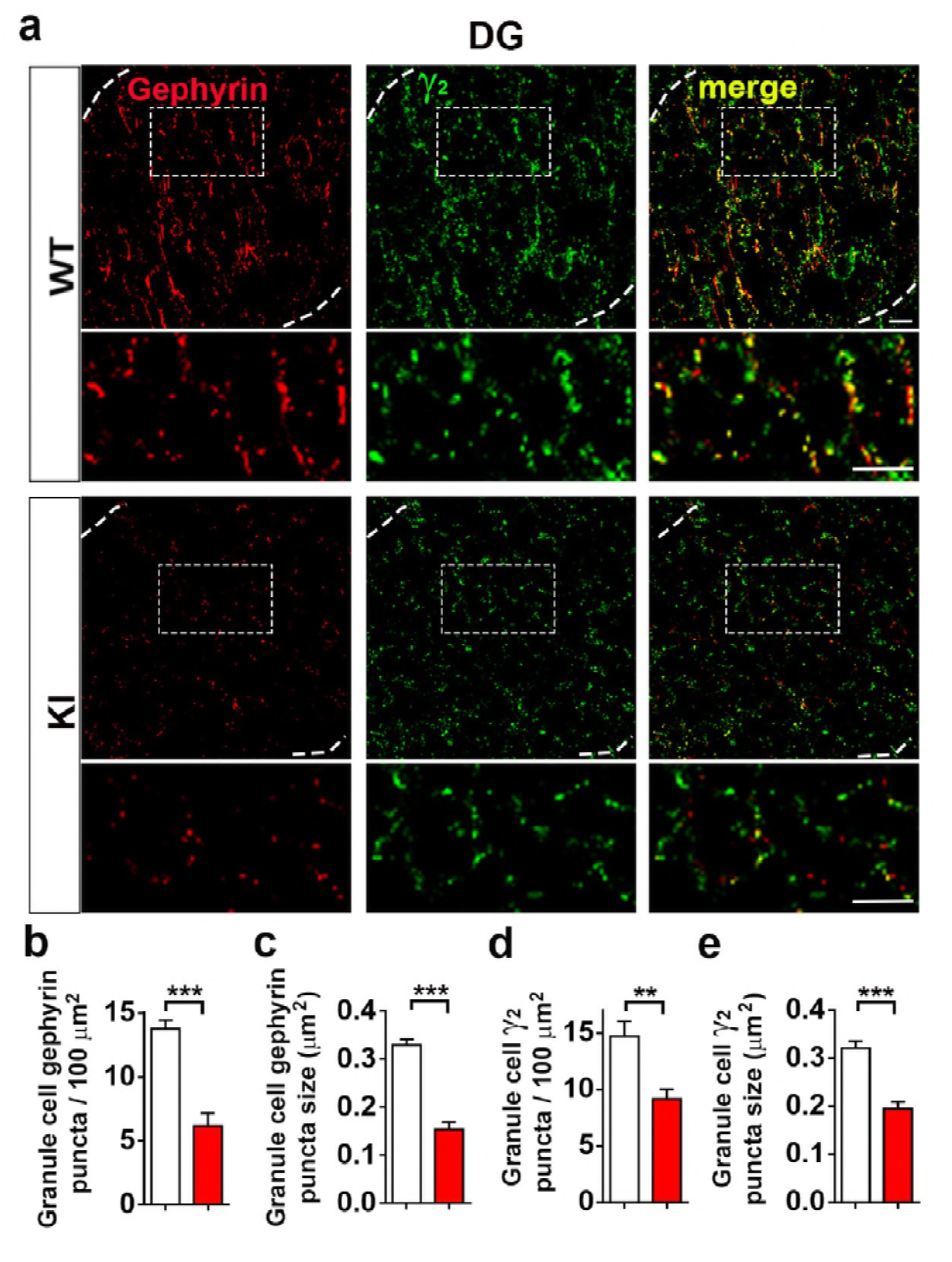
Reduced GABAergic postsynaptic components in the hippocampus of NL2 R215H KI mice. **(a)** Representative images of gephyrin, GABA_A_ receptor γ2 subunit and merged immunostaining in the granule cell layer of WT and R215H KI mice. **(b-c)** Quantification of gephyrin puncta number and size at granule cell soma region. **(d-e)** Quantification of γ2 puncta number and size at the same region as gephyrin. WT = 9 slices from 2 mice, R215H KI = 12 slices from 2 mice. Scale bar = 10 μm. Student’s *t*-test was used for analysis and data were shown as Mean ± SEM, *P < 0.05, **P < 0.01, ***P < 0.001.

**Figure 3.**
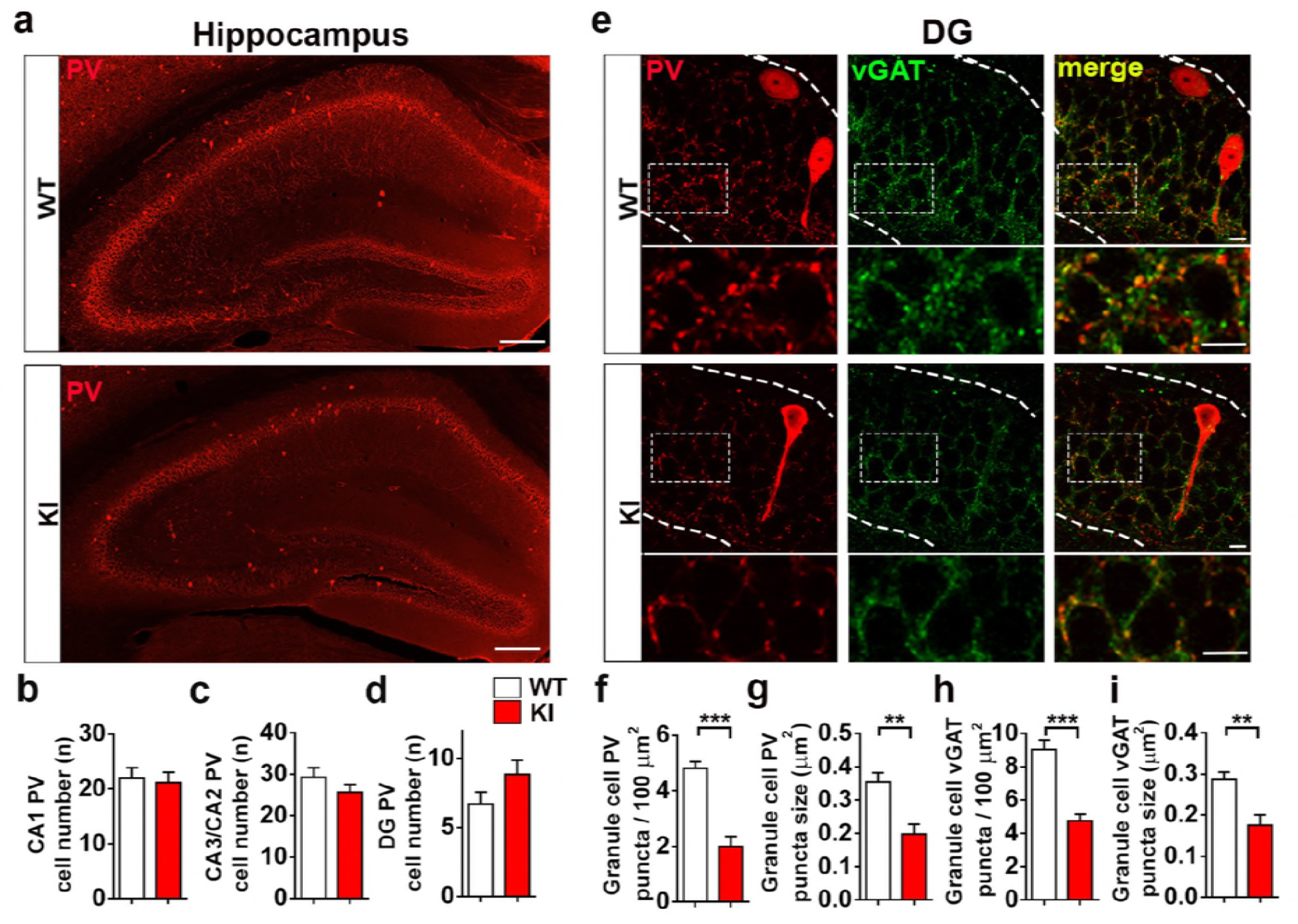
Reduced GABAergic presynaptic components in the hippocampus of NL2 R215H KI mice. **(a)** Representative images of PV staining at the hippocampus in WT and R215H KI mice. Scale bar = 200 μm. **(b-d)** Quantification of PV-positive neurons at DG, CA2/3, and CA1 region. WT, n= 14 slices / 5 mice; KI, n= 14 slices / 5 mice. **(e)** Representative images of PV, vGAT and merged immunostaining in the granule cell layer of WT and R215H KI mice. **(f-g)** Quantification of PV puncta number and size that targeted to the granule cell layer. **(h-i)** Quantification of vGAT puncta number and size that targeted to the same region. WT = 12 slices / 5 mice, R215H KI = 12 slices / 5 mice. Scale bar = 10 μm. Student’s *t* test was used for analysis and data were shown as Mean ± SEM, *P < 0.05, **P < 0.01, ***P < 0.001.

### Impaired GABAergic neurotransmission in NL2 R215H KI mice

We next investigated the function of inhibitory neurotransmission in the R215H KI mice. Whole-cell patch-clamp recordings were performed on dentate granule cells in acute brain slices of adult WT and homozygous R215H KI mice. We found that both the frequency and amplitude of miniature inhibitory postsynaptic currents (mIPSCs) were significantly decreased in the granule cells of R215H KI mice (Figure 4a-d; Frequency: WT = 8.28 ± 2.21 Hz, KI = 3.98 ± 0.78 Hz, p = 0.041; Median amplitude: WT = 41.3 ± 2.9 pA, KI = 32.7 ± 1.8 pA, p = 0.019; Student’s t-test). In contrast, there was no significant change of miniature excitatory postsynaptic currents (mEPSCs) in the dentate granule cells of R215H KI mice compared to WT mice (Figure 4e-h), indicating that NL2 R215H mutation is primarily affecting inhibitory neurotransmission.

**Figure 4.**
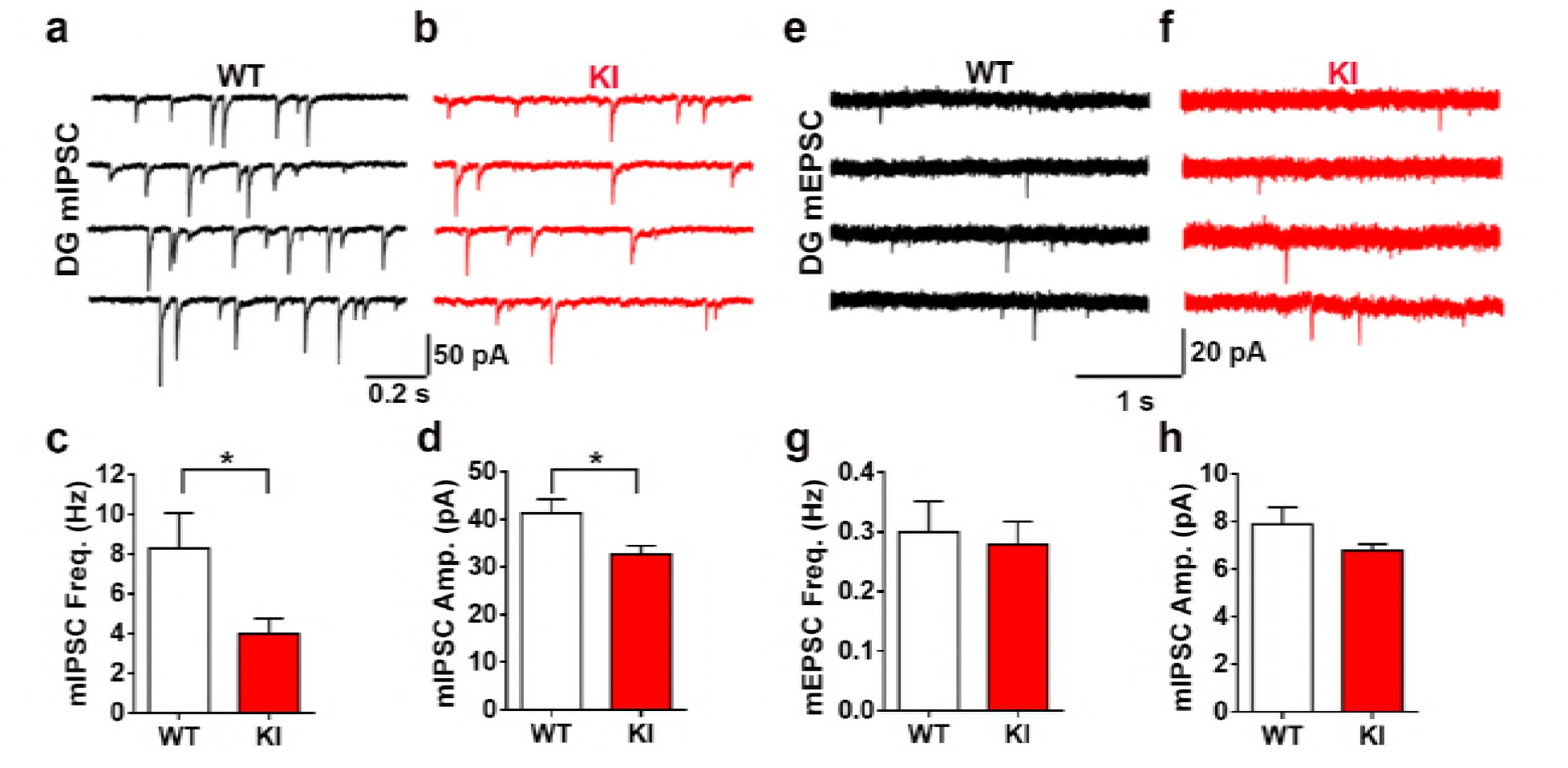
NL2 R215H KI mice have decreased inhibitory synaptic transmission at the hippocampal region. **(a-b)** Representative traces of miniature inhibitory postsynaptic currents (mIPSCs) recorded from DG granule cells in hippocampal slices of WT (black) and R215H KI (red) mice. WT, n = 14 cells / 4 mice; R215H KI, n = 18 cells / 4 mice. (c-d) Quantification of the mIPSC frequency and amplitude (Student’s t-test). **(e-f)** Representative traces of miniature excitatory postsynaptic currents (mEPSCs) in the DG region of hippocampal slices from WT (black) and R215H KI (red) mice. WT, n = 11 cells / 4 mice; KI, n = 12 cells / 3 mice. **(g-h)** Quantification of the mEPSC frequency and amplitude (Student’s t-test). Data represent mean ± SEM; *P < 0.05, **P < 0.01, ***P < 0.001.

### Behavioral deficits in NL2 R215H KI mice

The significant reduction of inhibitory neurotransmission in the NL2 R215H mutant mice prompted us to further investigate whether such severe GABAergic deficits will result in any behavioral deficits. We first performed open field test and elevated plus maze test to measure the mouse anxiety level. In the open field test, we found that the KI mice spent significantly less time in the center region, although the total distance traveled was similar to the WT mice (Figure 5a-d). Consistently, in the elevated plus maze test, the KI mice spent much less time in the open arm compared to the WT mice, while the total travel distance was also similar between the KI and WT mice (Figure 5e-h). These results suggest that the R215H KI mice display an increased level of anxiety while their locomotion activity is relatively normal.

**Figure 5.**
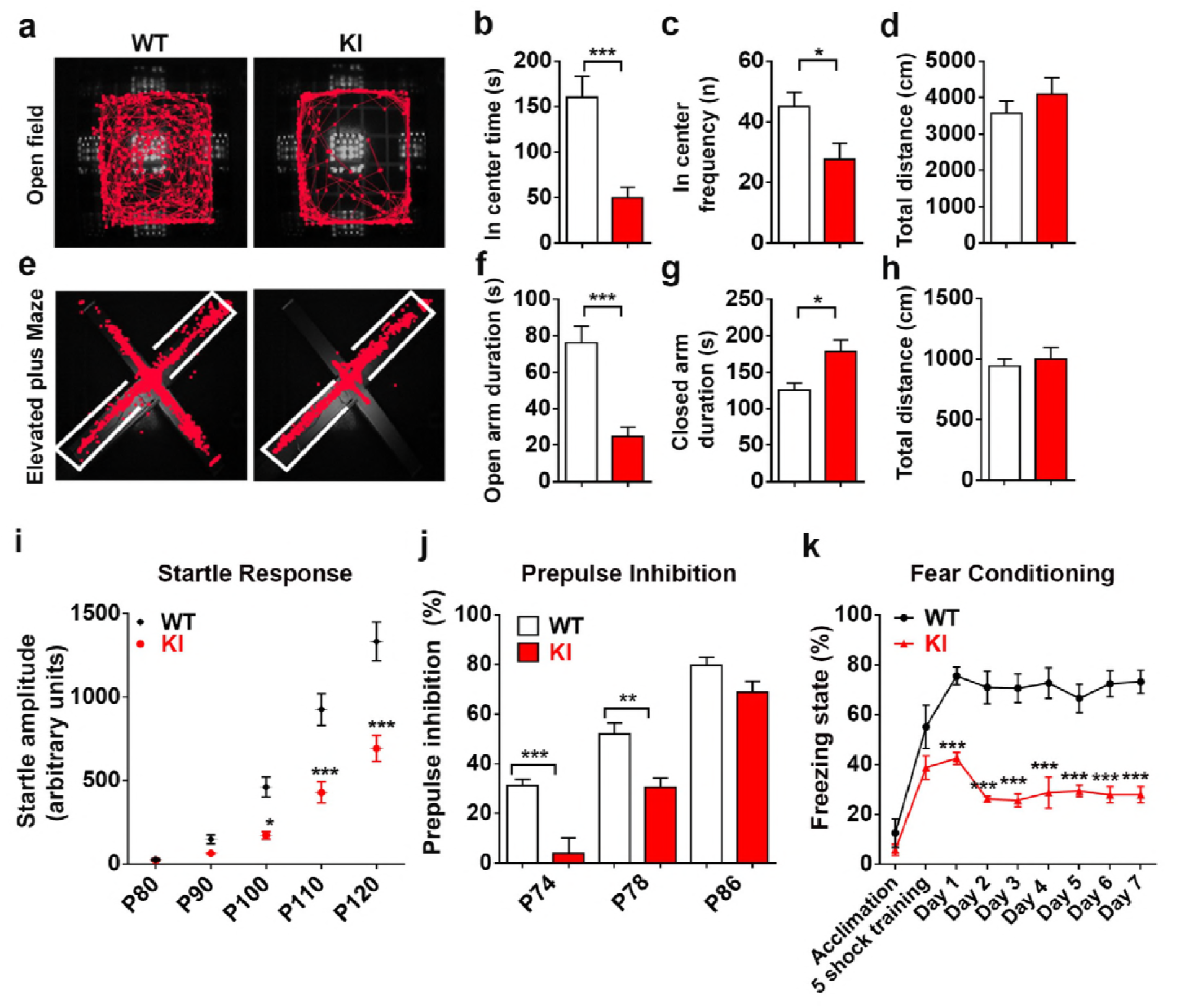
NL2 R215H KI mice display schizophrenia-like behaviors. **(a)** Representative running track of WT and R215H KI mice (male) in an open field within 10 minutes duration. **(b)** The center time of WT and KI mice spent in the open filed. **(c)** The frequency of WT and KI mice entering the center zone of the open field. **(d)** The total distance of WT and KI mice traveled in the open filed test. **(e)** Representative running track of WT and R215H KI mice (male) in elevated plus maze for 5 minutes. White line indicates closed-arms. **(f)** The quantified time spent in the open-arms of WT and KI mice. **(g)** The time spent in the closed-arms of WT and KI mice. **(h)** The total distance traveled in the elevated plus maze test. **(a-h)** WT mice n = 11, KI mice n = 12, age 2 to 3 months; Student’s t-test was used for analysis. **(i)** Startle response of WT and R215H KI mice (male) toward 80, 90, 100, 110, and 120 dB sound pulses. **(j)** The percentage of pre-pulse inhibition (PPI) to a pre-pulse of 74 dB, 78 dB, and 86 dB. WT mice n = 12, KI mice n = 9, age 3.5 months. Two-way ANOVA with Sidak’s multiple comparison test was used for analysis. **(k)** Contextual fear conditioning test. R215H KI mice exhibit significant reduction of freezing time when placed back in the test chamber after 1-7 days of shock training (Two-way ANOVA with Sidak’s multiple comparison test, genotype F_(1, 99)_ = 172.7, P < 0.0001, WT n = 8, KI n = 5, age 2 to 3 months).

We next examined in R215H KI mice the acoustic startle response and pre-pulse inhibition, a standard test for the sensory motor gating function often assessed in schizophrenia patients (Braff et al., 1992). R215H KI mice showed a significant reduction in the startle response when stimulated at 100 – 120 dB (Figure 5i). Furthermore, the pre-pulse inhibition was significantly impaired in the KI mice compared to the WT mice (Figure 5j). Together, these deficits of R215H KI mice suggest that this new transgenic mouse model may have symptoms of schizophrenia-like behavior.

To further characterize the R215H KI mice, we investigated their cognitive functions by conducting contextual fear conditioning test, a hippocampal dependent fear-learning test. We found that while the KI mice were capable to associate the conditioning chamber with foot-shock in the initial training, indicated by an increase of freezing state after foot-shock, they failed to retain the fear context memory in the following days when tested (Figure 5k), indicating an impaired cognition function. Importantly, the behavioral data shown above was all obtained from male mice, the female mice were also tested and exhibited the same trend (Figure S4).

### Impaired hippocampal activation toward acute stress in NL2 R215H KI mice

Schizophrenia is associated with abnormal response to stress (Walker and Diforio, 1997). Stress is known to activate the hypothalamic-pituitary-adrenal axis (HPA axis) and induce the hormone release of corticosterone (CORT) into circulation (Koob, 1999; McGill et al., 2006). To investigate the stress response of R215H KI mice, we put the WT and R215H KI mice into restraining tubes for one hour as an acute stress test. We found that R215H KI mice struggled much more intensively for a long time and excreted much more than the WT mice during the restraining test. After restraining, KI mice were more dirty and stinky than the WT mice (Figure 6a). In accordance, R215H KI mice showed a much higher level of CORT (384 ± 53 ng/ml) after restraining compared to the WT mice (215 ± 20 ng/ml). The baseline level of CORT was similar between WT (49 ± 4 ng/ml) and KI mice (36 ± 4 ng/ml) (Figure 6b; p = 0.0035 after restraint, Two way ANOVA followed with Sidak’s post hoc test). These results suggest that R215H KI mice have hyperactive HPA response toward stress.

**Figure 6.**
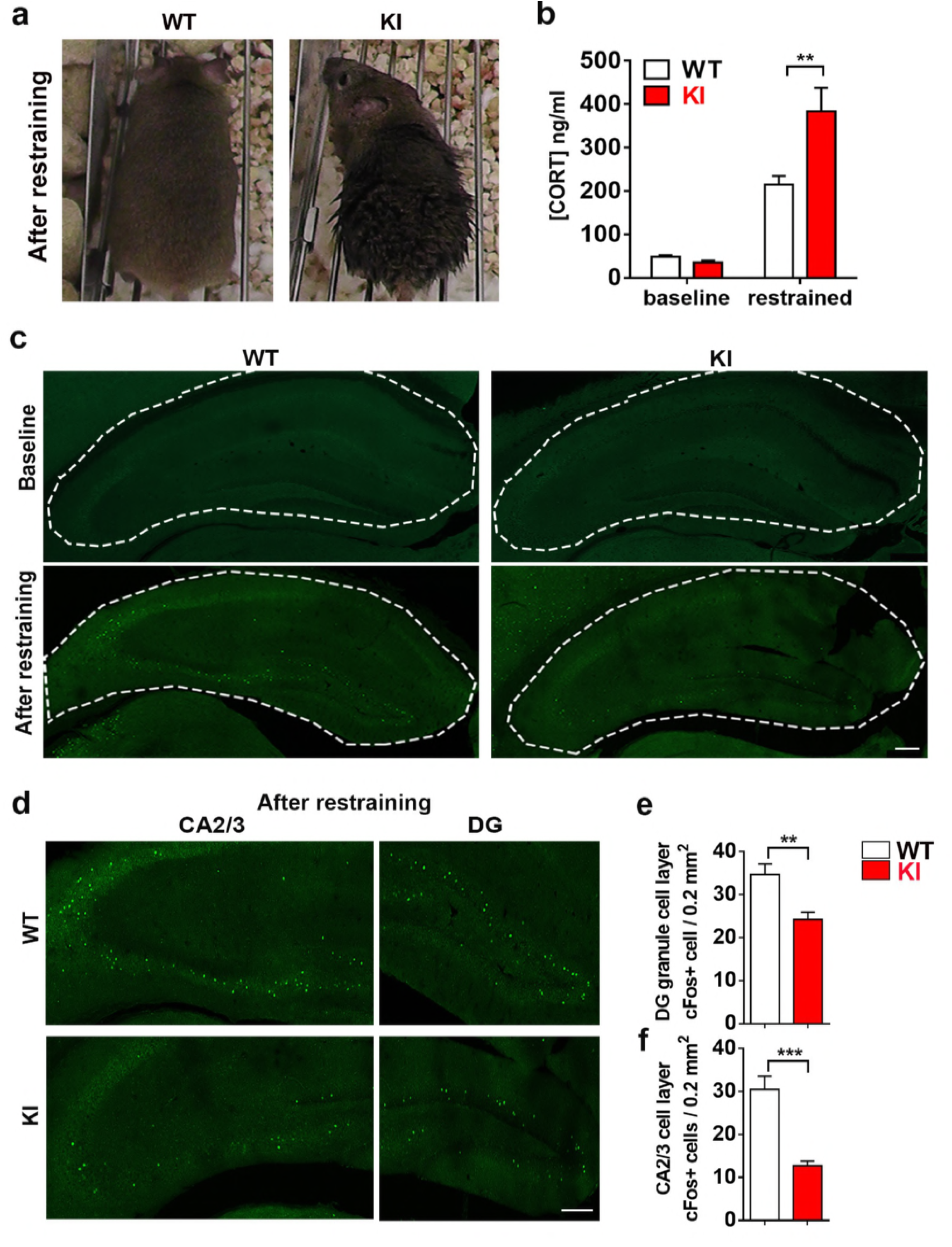
Hippocampal neurons have impaired activation toward acute stress in NL2 R215H KI mice. **(a)** Typical appearance of WT and KI mice after restraining. **(b)** Quantified corticosteroid level of WT and KI mice at the baseline level and after 1 h restraining. 4 to 8 mice were used for each genotype at each condition, age 4 to 6 months. Two-way ANOVA with Sidak’s multiple comparison test was used for analysis. **(c)** Upper row: representative images of baseline cFos immunoreactivity of WT and R215H KI hippocampus; bottom row: cFos immunoreactivity of WT and R215H KI hippocampus after restraining. **(d)** enlarged DG and CA2/3 region of WT and R215H mice after restraining. **(e)** Quantification of cFos-positive cells at DG granule cell layers. **(f)** Quantification of cFos positive cells at CA2/3 pyramidal cell layers. WT: n = 16 brain slices / 4 mice; R215H KI: n = 20 brain slices / 5 mice; scale bar = 200 μm. Student’s *t*-test was used for analysis. Data represent mean ± SEM; *P < 0.05, **P < 0.01, ***P < 0.001.

Following the activation of HPA axis, hippocampus will be activated as a negative feedback regulator and control the CORT level within normal range (Herman et al., 2012; Ulrich-Lai and Herman, 2009). To test whether hippocampus in R215H KI mice were activated following the acute stress, we used a naïve cohort of mice to perform the restraining test again. R215H KI and WT mice were subjected to restraint for half an hour and then sacrificed after 2 hours. Hippocampal activation was examined by assessing the expression level of an immediate early gene cFos (Morgan et al., 1987; Ramirez et al., 2013). At the baseline level, very few cFos-positive neurons were detected in the hippocampal regions in both WT and KI mice (Figure 6c, top row). After stress, we observed a significant increase of cFos-positive cells in the DG and CA2/3 regions of the hippocampus in WT mice (Figure 6c, bottom left). In contrast, the R215H KI mice showed much reduced cFos-positive cells in the same regions of hippocampus (Figure 6c, bottom right). This is better illustrated in the enlarged images showing the CA2/3 and DG regions of WT mice (Figure 6d, top row) and KI mice (Figure 6d, bottom row). Quantitative analysis confirmed the reduction of cFos-positive cells in both CA2/3 (Figure 6e) and DG (Figure 6f) regions in the KI mice. These results suggest that NL2 R215H KI mice had impaired hippocampal activation during acute stress.

## DISCUSSION

In the present study, we generated a unique mouse model carrying a single point mutation R215H of *NLGN2* gene that was originally identified from human schizophrenia patients. The NL2 R215H KI mice have impaired GABAergic synapse development, reduced inhibitory synaptic transmission, and decreased hippocampal activation in response to stress. Moreover, the R215H KI mice display anxiety-like behavior, impaired pre-pulse inhibition, cognitive deficits and abnormal stress response, partially recapitulating some of the core symptoms of schizophrenia patients. These results suggest that this newly generated R215H KI mouse line may provide a unique animal model for studying molecular mechanisms underlying schizophrenia and related neuropsychiatric disorders.

### GABAergic and behavioral deficits in NL2 R215H KI mice

NL2 plays important roles in regulating perisomatic GABAergic synapse development, phasic GABAergic transmission, and neural excitability (Babaev et al., 2016; Blundell et al., 2009; Chubykin et al., 2007; Gibson et al., 2009; Hines et al., 2008; Hoon et al., 2009; Jedlicka et al., 2010; Liang et al., 2015; Poulopoulos et al., 2009; Varoqueaux et al., 2004; Wohr et al., 2013). Consistent with our previous *in vitro* studies, the current *in vivo* work demonstrates that R215H mutation disrupts GABAergic synapse development. Functionally, NL2 R215H mutation caused a reduction of both frequency and amplitude of inhibitory neurotransmission. These results suggest that the R215H KI mice display more severe GABAergic deficits than the NL2 KO mice (Babaev et al., 2016; Chubykin et al., 2007; Gibson et al., 2009; Jedlicka et al., 2010; Poulopoulos et al., 2009). The more severe GABAergic deficits in our R215H KI mice might explain why they display more severe behavioral deficits than NL2 KO mice, such as PPI impairment, cognitive deficits, and abnormal stress response. Coincidentally, previous studies reported that NL3 R451C KI mouse also displayed stronger phenotypes than the NL3 KO mice (Etherton et al., 2011; Foldy et al., 2013; Tabuchi et al., 2007; Zhang et al., 2016). These evidences suggest that genetic mouse models based on mutations identified from patients may be more suitable than the germline KO mouse models for studying pathologic mechanisms of human diseases.

Behaviorally, NL2 R215H KI mice display an anxiety phenotype, which may be the result of decreased GABAergic inhibition (Blundell et al., 2009; Dalvi and Rodgers, 1996; Zarrindast et al., 2001). Interestingly, R215H KI mice also show impaired startle responses and deficits in pre-pulse inhibition (PPI). Previous study in rats has reported that disturbance of PV neuron development in the hippocampal DG region may cause reduction of PPI (Guo et al., 2013). A recent study also demonstrates that specific inhibition of PV neurons in the ventral hippocampus results in a reduction of both startle response and PPI (Nguyen et al., 2014). Consistent with these findings, we demonstrate here that our R215H KI mice display a significant reduction of PV innervation in the hippocampus, which may underlie the deficits of PPI. In contrast, the NL2 KO mice lack PPI deficit, which might be related to an insufficient loss of PV innervation at hippocampal regions (Wohr et al., 2013).

Another interesting observation is that the R215H KI mice are hyperactive after acute stress and are associated with impaired hippocampal activation. It has been reported that robust neuron activation requires low background activity before stimulus (Koistinaho et al., 1993; Rao et al., 2006). However, due to the reduction of GABAergic inhibition in our R215H KI mice, the baseline activity of hippocampal neurons may be elevated and thus a further activation of hippocampus by external stimulation will be dampened. The impaired activation of hippocampal neurons in R215H KI mice may contribute to the abnormal stress response we observed, as hippocampus acts like a “brake” during acute stress to prevent HPA axis from over activation (Hariri, 2015).

### NL2 R215H mutation and schizophrenia

It is well documented that schizophrenia patients show impaired pre-pulse inhibition as an abnormal sensorimotor gating deficit (Braff et al., 1992; Grillon et al., 1992). Many patients also have emotional symptoms such as anxiety and depression (Lewis and Lieberman, 2000). Additionally, patients are hypersensitive toward stress and certain patients have been found with altered HPA axis function (Bradley and Dinan, 2010). Intriguingly, R215H KI mice recapitulated these SCZ-like behaviors, suggesting a potential role of NL2 R215H in the development of schizophrenia symptoms. Furthermore, reduction of PV expression and PV-positive synapses is a prominent phenotype observed in SCZ patients (Lewis et al., 2001; Lewis et al., 2012; Lewis et al., 2005; Woo et al., 1998). The R215H mutation KI mice also show a significant reduction of PV innervation, consistent with the pathogenic deficit of SCZ patients. These GABAergic deficits, together with cognition and PPI deficits manifested in the KI mice, support the hypothesis that GABA dysfunction makes an important contribution to the cognitive and attention deficits of SCZ. Taken together, NLGN2 R215H single point mutation has a significant impact on GABAergic synapse development and the pathogenesis of neuropsychiatric disorders. Our newly generated NL2 R215H KI mice may provide a useful mouse model for the study of molecular mechanisms and drug development of neuropsychiatric disorders including schizophrenia.

## Materials and Methods

### NL2 R215H knock-in mice

The NL2 R215H knock-in mice were generated by homologous recombination in embryonic stem cells by Dr. Siu-Pok Yee’s team at the University of Connecticut Health Center. The detailed procedures are described in the *SI materials and methods.*

All the experimental mice were group housed (2-3 mice per cage) in home cages and lived at a constant 25 °C in a 12h light/dark cycle. Mice were given *ad libitum* access to food and water. Littermate or age and gender matched mice were used for experiments. All animal care and experiments followed the Penn State University IACUC protocol and NIH guidelines.

### Biochemical measurements

Protein levels were quantified using total brain homogenates from 3 groups of adult male littermates-WT, heterozygous and homozygous. The western blot system used was the standard Bio-Rad mini protein electrophoresis system and the procedure followed the system manual. LiCOR Odyssey Clx was used for protein signal detection. The antibodies used were Rb anti-Neuroligin 2 (1:1000, SYSY 129202), Rb anti-GAPDH (1:10000, Sigma G9545), and Gt anti-Rb 800 (1:15000, P/N 925-32210, P/N 925-32211). Detailed procedures are described in the *SI materials and methods.*

### Immunohistochemistry, Image Acquisition, and Image Analysis

Mouse brain slices were prepared at 20-40 μM and reacted with the primary antibodies Rb anti-Neuroligin 2 (1:1000, SYSY129203), Ms anti-Parvalbumin (1:1000, MAB1572), GP anti-vGAT (1:1000, SYSY 131004), Gephyrin (1:1000, SYSY 147011), GABAaR γ2 (1:1000 SYSY 224003), and c-Fos (1:5000 Sigma F7799). The fluorescent secondary antibodies used were Gt anti-Rb 488, Gt anti-Ms Cy3, and Gt anti-GP 647. Images were taken with the Olympus FV1000 confocal microscope. The number of neurons and the density and size of synaptic puncta were analyzed with the NIH ImageJ software (NIH, Bethesda, MD, USA). A detailed description of the experimental procedures is in the *SI materials and methods.*

### Slice electrophysiology

Horizontal acute hippocampal slices were used for whole-cell patch clamp recordings. Miniature inhibitory or excitatory postsynaptic currents (mIPSCs or mEPSCs) were pharmacologically isolated by including DNQX and APV or picrotoxin together with tetrodotoxin in artificial cerebrospinal fluid. Details are in the *SI materials and methods.*

### Behavioral tests

Overview: The mice for behavior tests were group housed by genotype. All tests were performed during 1 pm to 6 pm. Three cohorts of mice were used: First cohort of mice was tested for open field and elevated plus maze at 2 to 3 month and tested for the startle response and prepulse inhibition test at 3.5 months. Second cohort of mice was used for contextual fear conditioning test at 2-3 month. Third cohort of mice was used for restraining and corticosteroid serum level test at 4 to 6 months. The open field test and elevated maze data were analyzed by Noldus Ethovision XT 8.0 software. PPI test and CORT test were analyzed with the researcher blind to genotype. Detailed procedures are in *SI materials and methods.*

## Acknowledgements

We would like to thank Dr. Thomas Fuchs for providing advices on behavioral tests, Yuting Bai for providing initial genotyping support. We thank all members from Chen lab for thoughtful suggestions. This study is supported by grants from NIH (MH092740 and MH083911) and Charles H. “Skip” Smith Brain Repair Endowment Fund to G.C.

## SUPPLEMENTAL INFORMATION

### A. DETAILED EXPERIMENTAL PROCEDURES

#### Generation of NL2 R215H KI mice

A 11.2 kb mouse genomic clone encompassing exons 3 to 8 of neuroligin 2 was retrieved from BAC clone RP24-65N11 and subcloned into pL253 using gap repair method (1). The point mutation G644A (R215H) was introduced into exon 4 by unique restriction site elimination (2). A single LoxP site was inserted into intron 7 approximately 350 bp 3’ downstream of exon 7 by recombineering as described by Liu et al (1), the second LoxP site together with Frt-PGKneo-Frt cassette in reverse orientation was inserted in intron 3 approximately 500 bp 5’ upstream of exon 4. The targeting vector was linearized by Notl digestion and then electroporated into ES cells derived from a F1 (129SvEv X C57BL6/J) embryo. ES cells were then cultured in the presence of G418 and Gancyclovir 48 hours after electroporation. Drug resistant clones were picked into 96-well plates. Genomic DNAs prepared from duplicated plates were used to identify targeted ES clones using nested long-range PCR. The 3’ arm was identified by nested long-range PCR using the primer pair, *Nlgn2* ScR1: 5’-CAACCCCTAGCCTTCCTTACC and 452ScF1: 5’-CTTCTGAGGCGGAAAGAACCA, followed by a second primer pair, *Nlgn2* ScR2: GAGGATGCGGATTCGGCTCCAG and 451ScF2: 5’-CGAAGTTATTAGGTGGATCC, amplifying a 4.56kb fragment. The 5’arm was screened by nested long range PCR using primer pair, Nlgn2 ScF1: 5’-GGTAACTTTACATGCAGGCC and 452ScR1: 5’-TTCCAGACTCCTCATGCCTA followed by a second primer pair, Nlgn2 ScF2: 5’-CCTGTACCTCCTCATTGTGTAC and 452ScR2: GGAATTGGGCTGCAGGAATTCC, amplifying a fragment of 4kb. Three independent targeted ES clones were injected into blastocysts of CD-1 mice to generate chimeric animals and 2 chimeric males derived from each ES clone were used for breeding with ROSA26-Flpe female (Jackson Laboratory stock No. 003946, the flip mice were backcrossed with C57BL6/J over 10 generations) to remove the PGKneo cassette.

Germline transmission was determined by Lox gtF and Lox gtR primer (Lox gtF: 5’-GCAACTTGGTCTGCATCTAAG; Lox gtR: 5’-GAAGGGGAGGGAGAAGTCTGA), This primer pair amplify a fragment of 459 bp specific to mutant allele (in LoxP site) and a fragment of 367 bp specific to wild type allele (in intron 7) (Fig S1a). Genotyping was also confirmed by another primary pair Frt grF (5’-CAGCCATCTAAAGGATTCTTTG) and Frt gtR (5’-GGGCTGGAGGTGCTGGGAGG). The NL-2 R215H mutation was confirmed by sequencing PCR product of I3F primer (5’GTCCCCGATCTCCCGGCCCACC) and E5R primer (5’-GCAGCCTGGTCACCAGTGCTGAG) amplifying the *Nlgn2* exon 4. I3F primer was used for sequencing. PCR product were purified by QIAquick PCR purification Kit (Cat#28104) and sequenced by Penn State Genomics Core facility. The results were viewed and analyzed by sequencing scanner software 1.0 (Fig S1b).

NL2 R215H F1 heterozygotes (offspring of chimera mice X ROSA26-Flpe mice) were mated through non littermate heterozygotes X heterozygotes strategy. The mouse colony was maintained on a mixed genetic background. The wild type, NL2 R215H heterozygotes and NL-2 R215H homozygotes were weaned at postnatal 24 to 28 days.

#### Biochemical experiments

3 groups of male samples were used for analysis. Pierce IP lysis buffer were used for lysing mouse brain tissue (Thermo Scientific Prod# 87788), protease inhibitor cocktail (1: 200 Sigma P8340), phosphatase inhibitor (1: 200 Sigma P5726) and PMSF (1:100 Sigma P7626 10mg/ml) were added in the lysis buffer. Lysates were homogenate on ice using homogenate tube (wheaton 2ml, 7 ml). The sample then were centrifuged at 10,000g 4 °C for 10 minutes. The supernatant was collected and boiled with 4X loadingdye (NP0007, life technologies) and 1 % of β - Mercaptoethanol at 95 °C for 5 minutes. The sample were directly used for western blot or stored at -20 °C for further use.

#### Immunohistochemistry, Image Acquisition and Image Analysis

Mice were rapidly anesthetized and perfused with ice cold ACSF followed by 40 ml 4% PFA in 0.1M PB, post fixed in 4% PFA overnight in 4 °C. After fixation, brains were dehydrated by 30% sucrose in 0.1M PB. Slices were frozen on sliding microtome at -25 °C and sectioned at 30 um. For c-Fos staining, the mice were restrained for 0.5 hour and waited for 2 hours before perfusion. Brains were post-fixed for 48 hours before dehydration. Slices were sectioned at 40 um.

For gephyrin and GABA_A_ γ2 subunit, the brain slices were prepared following the method described by Schneider Gasser et. al.(3). Briefly, the brain slices were lively sectioned at 300 μm and post fixed for 30 min. After that, the brain slices were rinsed by 1xPBS and dehydrated in 30% sucrose for 3-5 hours. The brain slices were sectioned again at 20 μm in cryostat at -20 °C.

Slices were incubated in blocking solution for 1 hour (0.3% triton 5% NGS in 1X PBS), and then were incubated with primary antibody (dissolved in blocking solution) overnight in 4 °C. On second day, brain slices were washed 3 times with 1X PBS, and then incubated in secondary antibody (dissolved in 0.05% triton 5% NGS in 1X PBS) for 1.5 hours in room temperature. Slices were mounted with mounting solution (Invitrogen P36931). For c-Fos staining, the slices were incubated in 2% triton 10% NGS in PBS for 1 hour followed by primary antibody c-Fos (Sigma F7799) incubation (0.3% triton, 10% NGS in PBS) and secondary antibody incubation (0.05% triton, 10% NGS in PBS).

Synaptic puncta images were acquired by FV 1000 confocal microscope PlAPON 60X O NA 1.42 Objective lens at pixel size of 100 um and optical sections of 1 um. 3 sections were acquired from the subsurface of the brain slice for each image. Images were scanned by sequential line scanning at 8 us/pixel and the image resolution was 1024X1024 pixels with a data depth of 12 bits. Laser transmissivity and channel gain/offset parameters for each antibody were determined by wild type brain slices. All the images were acquired using identical image settings for laser power, channel gain/offset for a given immunostaining marker.

The image was analyzed by ImageJ software. The wild type group of images were analyzed first and a proper threshold were determined for just covering all the synaptic puncta signals. The same threshold was applied to all the images within the same set of immunostaining. Analyze particle function were used for puncta number quantification within granule cell layers or pyramidal cell layers.

#### Slice electrophysiology

For miniature IPSC and EPSC recording at DG region, adult hippocampal slices were prepared from 6 months old male or female littermates following protocol describe by Ting et al. and Zhao et al. (4, 5). NMDG (N-Methyl-D-glucamine) recovery method was slightly modified based on our own experimental condition. Adult mice were quickly anesthetized by 2.5% Avertin and perfused with oxygenated NMDG cutting solution (in mM): 93 NMDG, 93 HCl, 2.5 KCl, 1.2 NaH_2_PO4, 30 NaHCO3, 20 HEPES, 15 Glucose, 5 Sodium ascorbate, 2 Thiourea, 3 Sodium pyruvate, 10 MgSO4.7H_2_O, 0.5 CaCl_2_, 12 N-Acetyl-L-cysteine (PH 7.3-7.4 adjusted by HCl, Osm 300-310), the brains were quickly taken out and cut in the oxygenated NMDG cutting solution at room temperature using Leica 1200S. The 300 um brain slices were recovered at 33.0 ± 0.5 °C in oxygenated NMDG solution for 10-15 minutes. Brain slices were then transferred to the modified HEPES holding aCSF (in mM): 92 NaCl, 2.5 KCl, 1.2 NaH_2_PO4, 30 NaHCO3, 20 HEPES, 15 Glucose, 5 Sodium ascorbate, 2 Thiourea, 3 Sodium pyruvate, 2 MgSO4.7H_2_O, 2 CaCl_2_, 12 N-Acetyl-L-cysteine (PH 7.3-7.4 adjusted by NaOH, Osm 300-310) until recording. Recorded slice was transferred to a submerged recording chamber where they were continuously perfused (3 ml/min) with normal aCSF (in mM: 124 NaCl, 2.5 KCl, 1.2 NaH_2_PO4, 24 NaHCO3, 5HEPES, 13 Glucose, 2 MgSO4.7H2O, 2 CaCl_2_, saturated by 95% O2/5% CO2 at 33 °C (TC-324B, Warner instruments Inc). Slices were visualized with infrared optics using an Olympus microscope equipped with DIC optics. For mIPSC recording, the brain slices were incubated in aCSF with 0.5 uM TTX, 50 uM AP5, 10 uM DNQX. The internal pipette solution used was (in mM) 120 CsCl, 5 NaCl, 1 MgCl2, 10 HEPES, 0.5 EGTA, 0.3 Na2GTP, 3 MgATP. For mEPSC recording, the slices were incubated in ACSF with 0.5 uM TTX and 50 uM picrotoxin. The internal pipette solution was 135 CsMeSO4, 8 NaCl, 0.2 EGTA, 10 HEPES, 4 MgATP, 0.3 Na_2_GTP, 5 Tris-phosphocreatine. The pippette resistance was 3~5 MΩ, access resistance of recorded cells was less than 30 MΩ. Data were collected with a MultiClamp 700A amplifier and pCLAMP9 software (Molecular Devices). The synaptic events were analyzed using Mini Analysis Program (Synaptosoft, New Jersey, USA). over 200 events or 2 minutes recording periods were analyzed for each cell. Events detecting threshold was set at 10 pA. The analysis results were further confirmed visually.

#### Behavioral tests

##### Open field test and Elevated plus maze test

Open field tests were conducted in a 50×50×30 cm^3^ open field arena which was placed in a dark room (with red light). (The camera has built-in infra-red detection). The mice were allowed to roam the enclosure for 10 minutes under video monitoring. The center area was 20×20 cm^2^. The elevated plus maze was 25 cm (open arms) × 25 cm (closed arms) × 30 cm (height) and was placed in the same dark room as the open field test. Dim light (white LED light, 203 mA) was shone upon the maze during the test. The mice were allowed to explore the maze for 5 minutes under video tracking. The number of mice in the cohort: male WT n = 11, male KI n = 12; female WT n = 9, female KI n =10.

#### Pre-pulse inhibition test

The mice were subjected to the startle box once per day for 3 days before the formal test (5-10 minutes per session) in order to adapt them to the experimental environment. For the sound test, 8 trials of pulse at 80 dB (p80 dB), p90 dB, p100 dB, p110 dB, and p120 dB (each pulse for 40 ms) were randomly arranged and given to tested mice with a background noise of 70 dB (SR-Lab San Diego Instruments). The interval time between each trial was randomly set between 10 to 20 seconds by the program. The startle amplitude of the mice toward each pulse was averaged over the 8 trials. Outliers were excluded by Average ± 2*STDEV.

For the pre-pulse inhibition test, the mice first underwent 4 trials of p120. Next, they were subjected to a random ordering of 8 trials of p120 and 8 trials of each prepulse p74 dB, p78 dB, p86 dB (each pre-pulse was 20 ms followed by 100 ms of waiting and 40 ms of p120; the startle amplitude was recorded during the p120 window). The interval time between each trial were randomly generated by the program between 10 and 20 seconds. Four trials of p120 dB were given at the end of test. The total program was 25 minutes. The inhibition percentage was calculated by the formula (p120-pp) /p120*100. Outliers were excluded by Average ± 2* STDEV. The number of mice in the cohort: male WT n = 12, male KI n = 9; female WT n = 7, female KI n= 13.

#### Contextual fear conditioning

Mice were placed in the conditioned chamber (white noise on 55 dB) for 2 minutes before the unconditioned stimulus (foot shock; 2 ms, 0.2 mA) to explore the environment and remember the context. During training, mice were given 5 times of footshock with 1 minute interval. To measure the memory retention of contextual fear conditioning, mice were placed back in the test chamber 1 day after training for 7 days. Male WT n = 8, KI n = 5; Female WT n = 6, KI n = 5.

#### Corticosterone test

For the baseline corticosterone (CORT) test, experiments were conducted between 3-5 PM (1-3 h before the light turned off), the mice were rapidly anesthetized with isoflurane and decapitated (within 2 min of removing them from their cage). Blood was collected in a BD microcontainer (Cat#365956) and centrifuged for 8 minutes at 8000 rpm. Serum was then collected and stored at -80 °C until use. For the restraining-induced CORT test, mice were anesthetized and decapitated immediately following a 1-hour period of restraint. The blood sample collection procedure was the same as the baseline sample collection. The experiments were conducted within the same time frame as the baseline test. Four to eight mice were used for each genotype at both baseline and restraint conditions.

## SUPPLEMENTARY FIGURES and FIGURE LEGENDS

**Figure S1.**
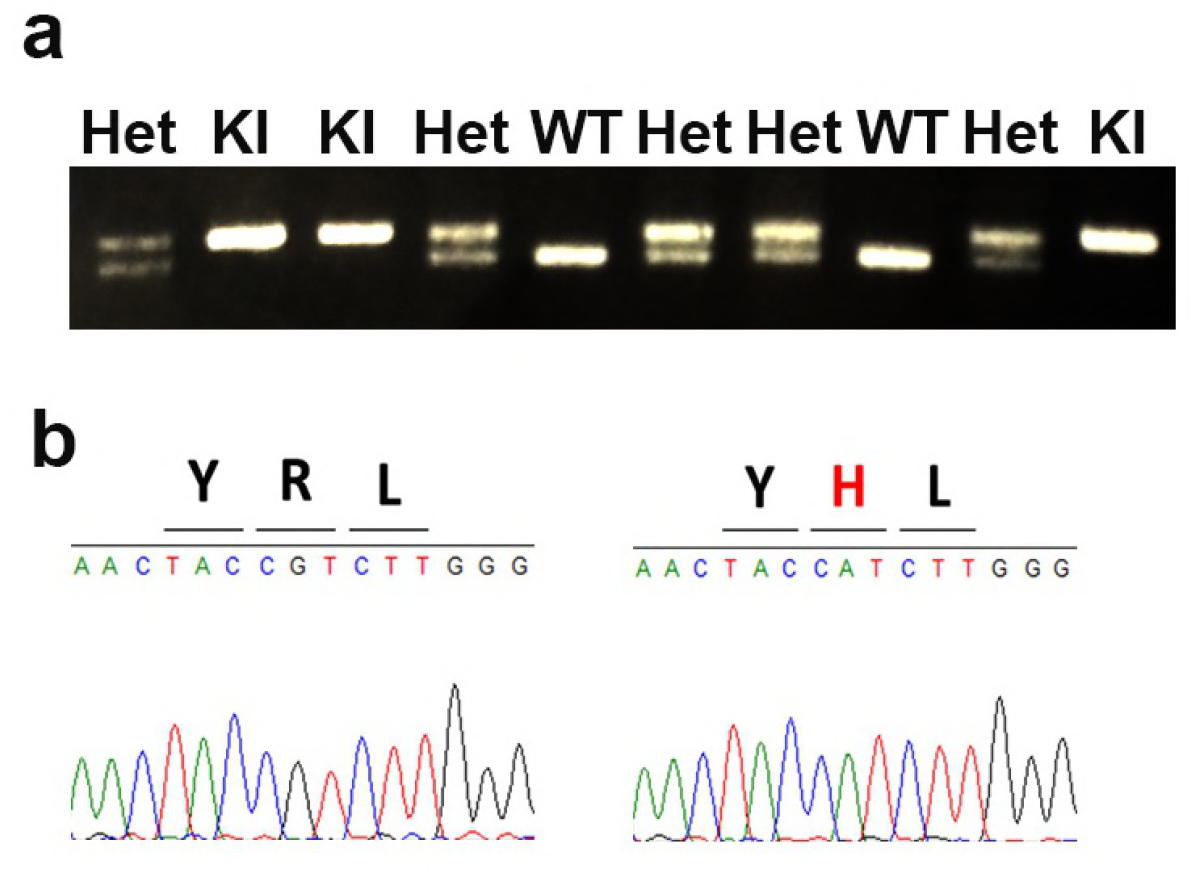
Generation of NL2 R215H KI mice. **(a)** A representative litter of offspring from R215H Heterozygous mating. Mouse genotype was determined by LoxgtR and LoxgtF primer. **(b)** Identification of NL2 R215H in mouse genome. Left: sequencing of WT *Nlgn2* exon 4 around G644 site. Right: sequencing of NL2 R215H homozygote at the same region as WT, the CGT codon was switched to CAT codon.

**Figure S2.**
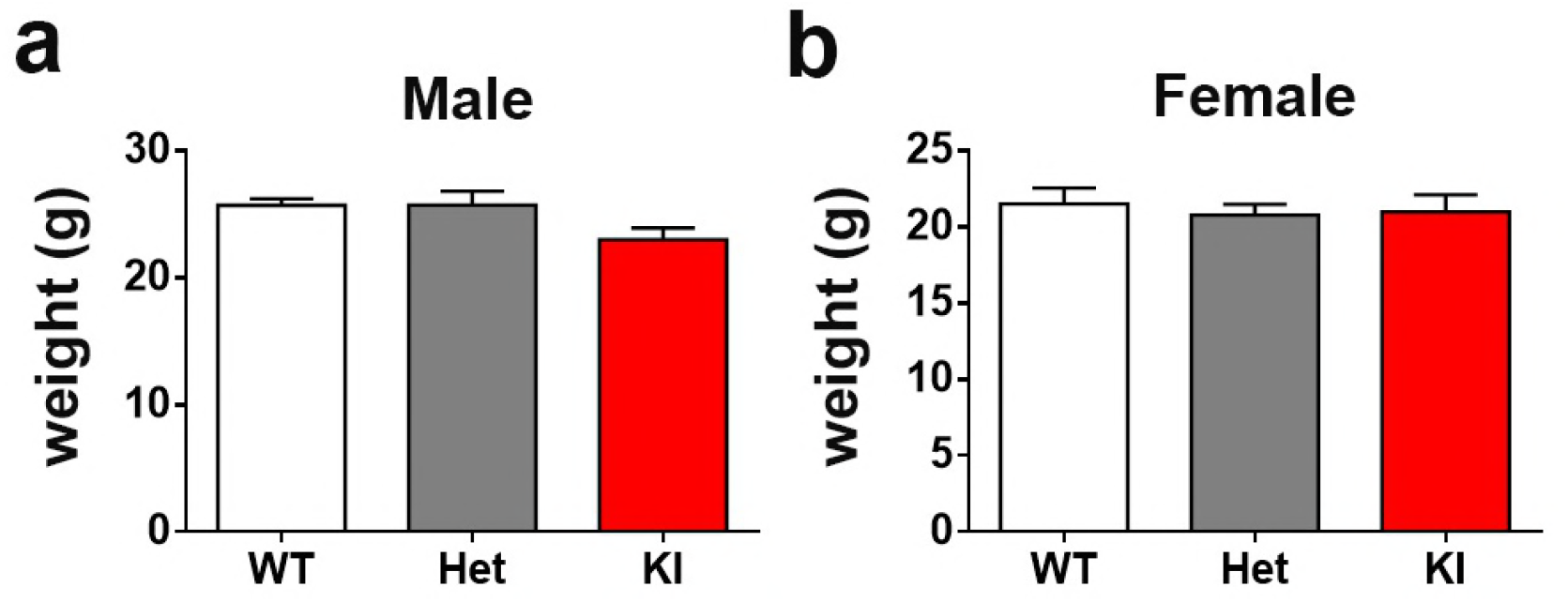
Body weight of WT, NL2 R215H Het, and NL2 R215H KI mice at 2 months age. **(a)** For male mice, WT = 11, Het = 14, KI = 7. **(b)** For Female mice, WT = 9, Het = 12, KI = 8. One way ANOVA test with Tukey multi-comparisons test was used for statistical analysis. Data were shown as Mean ± SEM.

**Figure S3.**
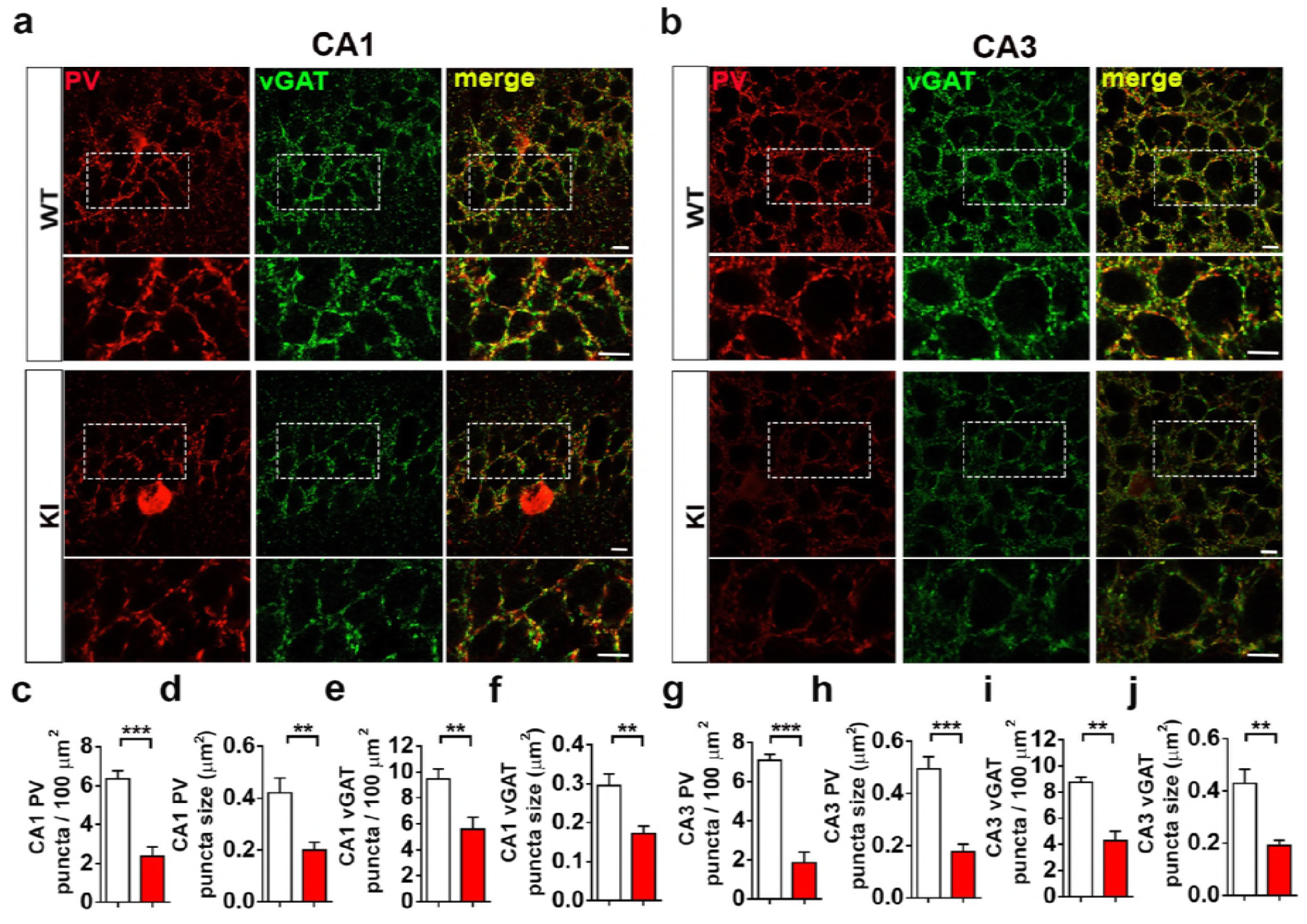
Parvalbumin and vGAT signal significantly reduced in NL2 R215H KI mice. **(a)** Representative image of PV and vGAT staining at CA1 in WT and R215H KI mice. Scale bar = 10 um. **(b)** Representative image of PV and vGAT staining at CA3 in WT and R215H KI mice. Scale bar = 10 um. **(c-f)** CA1 pyramidal cell soma region PV and vGAT puncta quantification. **(g-j)** CA3 cell soma region PV and vGAT puncta quantification. WT n = 9 slices / 3 mice; KI n = 7 slices / 3 mice; Student t tests were used for data analysis. Data were shown as Mean ± SEM. *P < 0.05, **P < 0.01, ***P < 0.001.

**Figure S4.**
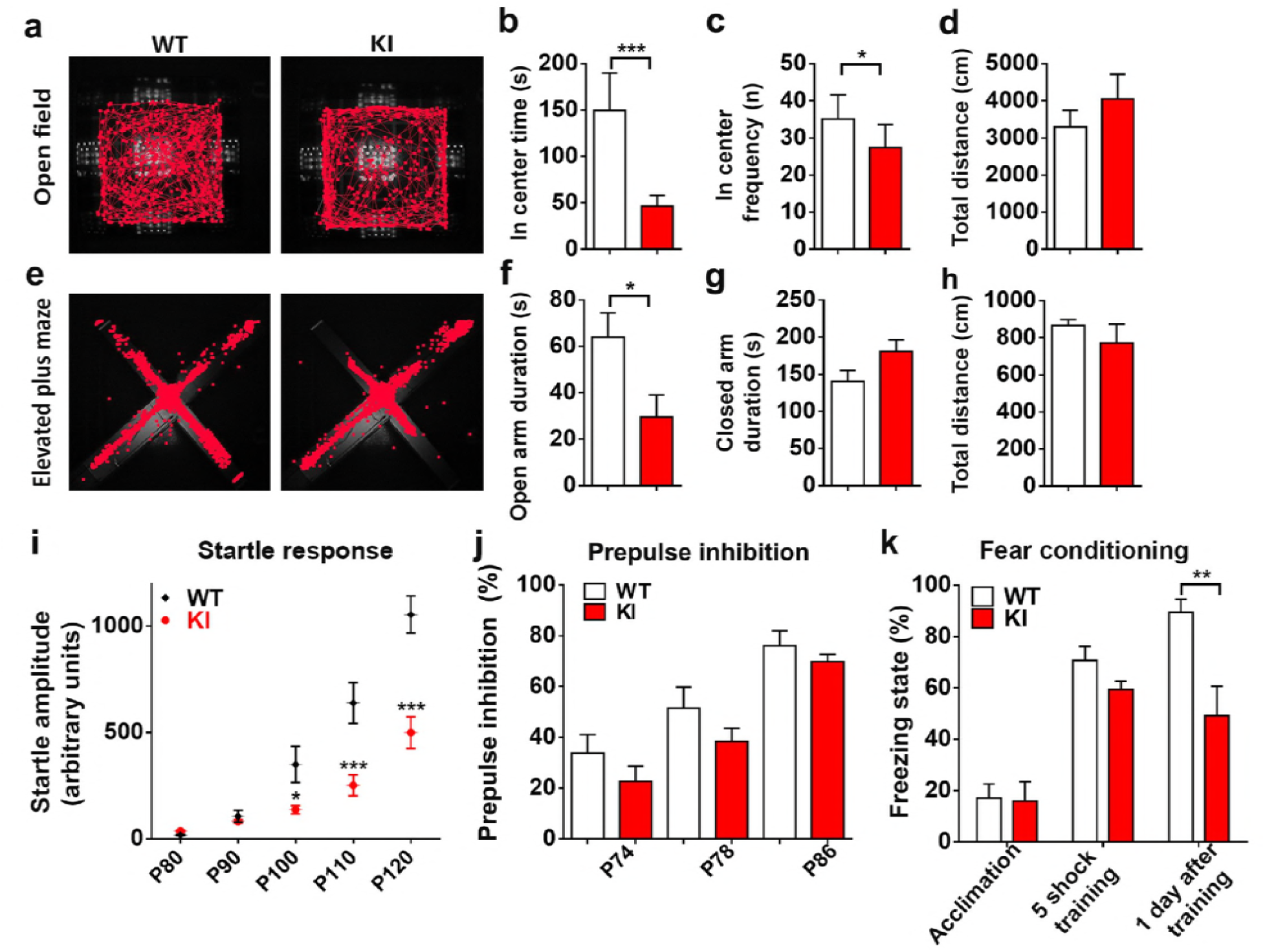
NL2 R215H female mice show schizophrenia-like behaviors. **(a)** Representative track of WT and R215H KI female mice running in open field within 10 minutes duration. **(b)** The center time (seconds) of WT and KI female mice spent in open filed within the trial. **(c)** The frequency of WT and KI female mice entering open field center zone. **(d)** The total distance of WT and KI female mice traveled in open field. **(e)** Representative track of WT and R215H KI female mice in elevated plus maze for 5 minutes. **(f)** The quantified time spent in open arms of WT and KI female mice. **(g)** The time spent in closed arms of WT and KI female mice. **(h)** The total distance traveled on elevated plus maze. **(a-d)** WT mice n=9, KI mice n=8; age at 3 months; **(e-h)** WT mice n = 9, KI mice n = 10, age 2 months. Student’s t tests were used for analysis. **(i)** Startle response of WT and R215H KI female mice toward 80, 90, 100, 110, and 120 dB sound pulses. **(j)** The percentage of prepulse inhibition of the startle response to a prepulse of 74 dB, 78 dB, and 86 dB. WT mice n=7, KI mice n=13, age 3.5 months. Two way ANOVA with Sidak’s multiple comparison test was used for analysis. (k) R215H KI female mice exhibited significant reduced freezing time 1 day after shock training. Twoway ANOVA with Sidak’s multiple comparison test, All data were shown as Mean ± SEM, *P < 0.05, **P < 0.01, ***P < 0.001.

## References

Babaev, O., Botta, P., Meyer, E., Muller, C., Ehrenreich, H., Brose, N., Luthi, A., and Krueger-Burg, D. (2016). Neuroligin 2 deletion alters inhibitory synapse function and anxiety-associated neuronal activation in the amygdala. Neuropharmacology 100, 56–65.

Baudouin, S.J., Gaudias, J., Gerharz, S., Hatstatt, L., Zhou, K., Punnakkal, P., Tanaka, K.F., Spooren, W., Hen, R., De Zeeuw, C.I., Vogt, K., and Scheiffele, P. (2012). Shared synaptic pathophysiology in syndromic and nonsyndromic rodent models of autism. Science 338, 128–132.

Blundell, J., Tabuchi, K., Bolliger, M.F., Blaiss, C.A., Brose, N., Liu, X., Sudhof, T.C., and Powell, C.M. (2009). Increased anxiety-like behavior in mice lacking the inhibitory synapse cell adhesion molecule neuroligin 2. Genes Brain Behav 8, 114–126.

Bradley, A.J., and Dinan, T.G. (2010). Review: A systematic review of hypothalamic-pituitary-adrenal axis function in schizophrenia: implications for mortality. Journal of Psychopharmacology 24, 91–118.

Braff, D.L., Grillon, C., and Geyer, M.A. (1992). Gating and habituation of the startle reflex in schizophrenic patients. Arch Gen Psychiatry 49, 206–215.

Bucan, M., Abrahams, B.S., Wang, K., Glessner, J.T., Herman, E.I., Sonnenblick, L.I., Alvarez Retuerto, A.I., Imielinski, M., Hadley, D., Bradfield, J.P., Kim, C., Gidaya, N.B., Lindquist, I., Hutman, T., Sigman, M., Kustanovich, V., Lajonchere, C.M., Singleton, A., Kim, J., Wassink, T.H., McMahon, W.M., Owley, T., Sweeney, J.A., Coon, H., Nurnberger, J.I., Li, M., Cantor, R.M., Minshew, N.J., Sutcliffe, J.S., Cook, E.H., Dawson, G., Buxbaum, J.D., Grant, S.F., Schellenberg, G.D., Geschwind, D.H., and Hakonarson, H. (2009). Genome-wide analyses of exonic copy number variants in a family-based study point to novel autism susceptibility genes. PLoS Genet 5, e1000536.

Budreck, E.C., and Scheiffele, P. (2007). Neuroligin-3 is a neuronal adhesion protein at GABAergic and glutamatergic synapses. Eur J Neurosci 26, 1738–1748.

Chih, B., Afridi, S.K., Clark, L., and Scheiffele, P. (2004). Disorder-associated mutations lead to functional inactivation of neuroligins. Hum Mol Genet 13, 1471–1477.

Chubykin, A.A., Atasoy, D., Etherton, M.R., Brose, N., Kavalali, E.T., Gibson, J.R., and Sudhof, T.C. (2007). Activity-dependent validation of excitatory versus inhibitory synapses by neuroligin-1 versus neuroligin-2. Neuron 54, 919–931.

Connor, S.A., Ammendrup-Johnsen, I., Chan, A.W., Kishimoto, Y., Murayama, C., Kurihara, N., Tada, A., Ge, Y., Lu, H., Yan, R., LeDue, J.M., Matsumoto, H., Kiyonari, H., Kirino, Y., Matsuzaki, F., Suzuki, T., Murphy, T.H., Wang, Y.T., Yamamoto, T., and Craig, A.M. (2016). Altered Cortical Dynamics and Cognitive Function upon Haploinsufficiency of the Autism-Linked Excitatory Synaptic Suppressor MDGA2. Neuron 91, 1052–1068.

Dalvi, A., and Rodgers, R.J. (1996). GABAergic influences on plus-maze behaviour in mice. Psychopharmacology 128, 380–397.

Dong, N., Qi, J., and Chen, G. (2007). Molecular reconstitution of functional GABAergic synapses with expression of neuroligin-2 and GABAA receptors. Mol Cell Neurosci 35, 14–23.

Durand, C.M., Betancur, C., Boeckers, T.M., Bockmann, J., Chaste, P., Fauchereau, F., Nygren, G., Rastam, M., Gillberg, I.C., Anckarsater, H., Sponheim, E., Goubran-Botros, H., Delorme, R., Chabane, N., Mouren-Simeoni, M.C., de Mas, P., Bieth, E., Roge, B., Heron, D., Burglen, L., Gillberg, C., Leboyer, M., and Bourgeron, T. (2007). Mutations in the gene encoding the synaptic scaffolding protein SHANK3 are associated with autism spectrum disorders. Nat Genet 39, 25–27.

Etherton, M., Foldy, C., Sharma, M., Tabuchi, K., Liu, X., Shamloo, M., Malenka, R.C., and Sudhof, T.C. (2011). Autism-linked neuroligin-3 R451C mutation differentially alters hippocampal and cortical synaptic function. Proc Natl Acad Sci U S A 108, 13764–13769.

Etherton, M.R., Blaiss, C.A., Powell, C.M., and Sudhof, T.C. (2009). Mouse neurexin-1alpha deletion causes correlated electrophysiological and behavioral changes consistent with cognitive impairments. Proc Natl Acad Sci U S A 106, 17998–18003.

Foldy, C., Malenka, R.C., and Sudhof, T.C. (2013). Autism-associated neuroligin-3 mutations commonly disrupt tonic endocannabinoid signaling. Neuron 78, 498–509.

Freedman, R. (2003). Schizophrenia. N Engl J Med 349, 1738–1749.

Gibson, J.R., Huber, K.M., and Sudhof, T.C. (2009). Neuroligin-2 deletion selectively decreases inhibitory synaptic transmission originating from fast-spiking but not from somatostatin-positive interneurons. J Neurosci 29, 13883–13897.

Grillon, C., Ameli, R., Charney, D.S., Krystal, J., and Braff, D. (1992). Startle gating deficits occur across prepulse intensities in schizophrenic patients. Biol Psychiatry 32, 939–943.

Guo, N., Yoshizaki, K., Kimura, R., Suto, F., Yanagawa, Y., and Osumi, N. (2013). A sensitive period for GABAergic interneurons in the dentate gyrus in modulating sensorimotor gating. J Neurosci 33, 6691–6704.

Hariri, A.R. (2015). Looking inside the disordered brain : an introduction to the functional neuroanatomy of psychopathology (Sunderland, Massachusetts, Sinauer Associates, Inc.).

Herman, J.P., McKlveen, J.M., Solomon, M.B., Carvalho-Netto, E., and Myers, B. (2012). Neural regulation of the stress response: glucocorticoid feedback mechanisms. Braz J Med Biol Res 45, 292–298.

Hines, R.M., Wu, L., Hines, D.J., Steenland, H., Mansour, S., Dahlhaus, R., Singaraja, R.R., Cao, X., Sammler, E., Hormuzdi, S.G., Zhuo, M., and El-Husseini, A. (2008). Synaptic imbalance, stereotypies, and impaired social interactions in mice with altered neuroligin 2 expression. J Neurosci 28, 6055–6067.

Hoon, M., Bauer, G., Fritschy, J.M., Moser, T., Falkenburger, B.H., and Varoqueaux, F. (2009). Neuroligin 2 controls the maturation of GABAergic synapses and information processing in the retina. J Neurosci 29, 8039–8050.

Ichtchenko, K., Hata, Y., Nguyen, T., Ullrich, B., Missler, M., Moomaw, C., and Sudhof, T.C. (1995). Neuroligin 1: a splice site-specific ligand for beta-neurexins. Cell 81, 435–443.

Insel, T.R. (2010). Rethinking schizophrenia. Nature 468, 187–193.

Jamain, S., Quach, H., Betancur, C., Rastam, M., Colineaux, C., Gillberg, I.C., Soderstrom, H., Giros, B., Leboyer, M., Gillberg, C., and Bourgeron, T. (2003). Mutations of the X-linked genes encoding neuroligins NLGN3 and NLGN4 are associated with autism. Nat Genet 34, 27–29.

Jamain, S., Radyushkin, K., Hammerschmidt, K., Granon, S., Boretius, S., Varoqueaux, F., Ramanantsoa, N., Gallego, J., Ronnenberg, A., Winter, D., Frahm, J., Fischer, J., Bourgeron, T., Ehrenreich, H., and Brose, N. (2008). Reduced social interaction and ultrasonic communication in a mouse model of monogenic heritable autism. Proc Natl Acad Sci U S A 105, 1710–1715.

Jedlicka, P., Hoon, M., Papadopoulos, T., Vlachos, A., Winkels, R., Poulopoulos, A., Betz, H., Deller, T., Brose, N., Varoqueaux, F., and Schwarzacher, S.W. (2010). Increased dentate gyrus excitability in neuroligin-2-deficient mice in vivo. Cereb Cortex 21, 357–367.

Kim, H.G., Kishikawa, S., Higgins, A.W., Seong, I.S., Donovan, D.J., Shen, Y., Lally, E., Weiss, L.A., Najm, J., Kutsche, K., Descartes, M., Holt, L., Braddock, S., Troxell, R., Kaplan, L., Volkmar, F., Klin, A., Tsatsanis, K., Harris, D.J., Noens, I., Pauls, D.L., Daly, M.J., MacDonald, M.E., Morton, C.C., Quade, B.J., and Gusella, J.F. (2008). Disruption of neurexin 1 associated with autism spectrum disorder. Am J Hum Genet 82, 199–207.

Kirov, G., Gumus, D., Chen, W., and Norton, N. (2008). Comparative genome hybridization suggests a role for NRXN1 and APBA2 in schizophrenia. Human molecular ….

Koistinaho, J., Hicks, K.J., and Sagar, S.M. (1993). Tetrodotoxin enhances light-induced c-fos gene expression in the rabbit retina. Brain Res Mol Brain Res 17, 179–183.

Koob, G.F. (1999). Corticotropin-releasing factor, norepinephrine, and stress. Biol Psychiatry 46, 1167–1180.

Levinson, J.N., Chery, N., Huang, K., Wong, T.P., Gerrow, K., Kang, R., Prange, O., Wang, Y.T., and El-Husseini, A. (2005). Neuroligins mediate excitatory and inhibitory synapse formation: involvement of PSD-95 and neurexin-1beta in neuroligin-induced synaptic specificity. J Biol Chem 280, 17312–17319.

Lewis, D.A., Cruz, D.A., Melchitzky, D.S., and Pierri, J.N. (2001). Lamina-specific deficits in parvalbumin-immunoreactive varicosities in the prefrontal cortex of subjects with schizophrenia: evidence for fewer projections from the thalamus. Am J Psychiatry 158, 1411–1422.

Lewis, D.A., Curley, A.A., Glausier, J.R., and Volk, D.W. (2012). Cortical parvalbumin interneurons and cognitive dysfunction in schizophrenia. Trends in Neurosciences 35, 57–67.

Lewis, D.A., Hashimoto, T., and Volk, D.W. (2005). Cortical inhibitory neurons and schizophrenia. Nature reviews Neuroscience 6, 312–324.

Lewis, D.A., and Lieberman, J.A. (2000). Catching up on schizophrenia: natural history and neurobiology. Neuron.

Liang, J., Xu, W., Hsu, Y.T., Yee, A.X., Chen, L., and Sudhof, T.C. (2015). Conditional neuroligin-2 knockout in adult medial prefrontal cortex links chronic changes in synaptic inhibition to cognitive impairments. Mol Psychiatry 20, 850–859.

McGill, B.E., Bundle, S.F., Yaylaoglu, M.B., Carson, J.P., Thaller, C., and Zoghbi, H.Y. (2006). Enhanced anxiety and stress-induced corticosterone release are associated with increased Crh expression in a mouse model of Rett syndrome. Proc Natl Acad Sci U S A 103, 18267–18272.

Morgan, J.I., Cohen, D.R., Hempstead, J.L., and Curran, T. (1987). Mapping patterns of c-fos expression in the central nervous system after seizure. Science 237, 192–197.

Nam, C.I., and Chen, L. (2005). Postsynaptic assembly induced by neurexin-neuroligin interaction and neurotransmitter. Proc Natl Acad Sci U S A 102, 6137–6142.

Nguyen, R., Morrissey, M.D., Mahadevan, V., Cajanding, J.D., Woodin, M.A., Yeomans, J.S., Takehara-Nishiuchi, K., and Kim, J.C. (2014). Parvalbumin and GAD65 interneuron inhibition in the ventral hippocampus induces distinct behavioral deficits relevant to schizophrenia. J Neurosci 34, 14948–14960.

Peça, J., Feliciano, C., Ting, J.T., Wang, W., Wells, M.F., Venkatraman, T.N., Lascola, C.D., Fu, Z., and Feng, G. (2011). Shank3 mutant mice display autistic-like behaviours and striatal dysfunction. Nature 472, 437–442.

Poulopoulos, A., Aramuni, G., Meyer, G., Soykan, T., Hoon, M., Papadopoulos, T., Zhang, M., Paarmann, I., Fuchs, C., Harvey, K., Jedlicka, P., Schwarzacher, S.W., Betz, H., Harvey, R.J., Brose, N., Zhang, W., and Varoqueaux, F. (2009). Neuroligin 2 drives postsynaptic assembly at perisomatic inhibitory synapses through gephyrin and collybistin. Neuron 63, 628–642.

Ramirez, S., Liu, X., Lin, P.A., Suh, J., Pignatelli, M., Redondo, R.L., Ryan, T.J., and Tonegawa, S. (2013). Creating a false memory in the hippocampus. Science 341, 387–391.

Rao, V.R., Pintchovski, S.A., Chin, J., Peebles, C.L., Mitra, S., and Finkbeiner, S. (2006). AMPA receptors regulate transcription of the plasticity-related immediate-early gene Arc. Nat Neurosci 9, 887–895.

Rothwell, P.E., Fuccillo, M.V., Maxeiner, S., Hayton, S.J., Gokce, O., Lim, B.K., Fowler, S.C., Malenka, R.C., and Sudhof, T.C. (2014). Autism-associated neuroligin-3 mutations commonly impair striatal circuits to boost repetitive behaviors. Cell 158, 198–212.

Scheiffele, P., Fan, J., Choih, J., Fetter, R., and Serafini, T. (2000). Neuroligin expressed in nonneuronal cells triggers presynaptic development in contacting axons. Cell 101, 657–669.

Schizophrenia Working Group of the Psychiatric Genomics, C. (2014). Biological insights from 108 schizophrenia-associated genetic loci. Nature 511, 421–427.

Song, J.Y., Ichtchenko, K., Sudhof, T.C., and Brose, N. (1999). Neuroligin 1 is a postsynaptic cell-adhesion molecule of excitatory synapses. Proc Natl Acad Sci U S A 96, 1100–1105.

Südhof, T.C. (2008). Neuroligins and neurexins link synaptic function to cognitive disease. Nature 455, 903–911.

Sullivan, P.F., Kendler, K.S., and Neale, M.C. (2003). Schizophrenia as a complex trait: evidence from a meta-analysis of twin studies. Archives of general psychiatry 60, 1187–1192.

Sun, C., Cheng, M.C., Qin, R., Liao, D.L., Chen, T.T., Koong, F.J., Chen, G., and Chen, C.H. (2011). Identification and functional characterization of rare mutations of the neuroligin-2 gene (NLGN2) associated with schizophrenia. Hum Mol Genet 20, 3042–3051.

Tabuchi, K., Blundell, J., Etherton, M.R., Hammer, R.E., Liu, X., Powell, C.M., and Sudhof, T.C. (2007). A neuroligin-3 mutation implicated in autism increases inhibitory synaptic transmission in mice. Science 318, 71–76.

Ulrich-Lai, Y.M., and Herman, J.P. (2009). Neural regulation of endocrine and autonomic stress responses. Nat Rev Neurosci 10, 397–409.

Varoqueaux, F., Jamain, S., and Brose, N. (2004). Neuroligin 2 is exclusively localized to inhibitory synapses. Eur J Cell Biol 83, 449–456.

Walker, E.F., and Diforio, D. (1997). Schizophrenia: a neural diathesis-stress model. Psychol Rev 104, 667–685.

Wohr, M., Silverman, J.L., Scattoni, M.L., Turner, S.M., Harris, M.J., Saxena, R., and Crawley, J.N. (2013). Developmental delays and reduced pup ultrasonic vocalizations but normal sociability in mice lacking the postsynaptic cell adhesion protein neuroligin2. Behav Brain Res 251, 50–64.

Woo, T.U., Whitehead, R.E., Melchitzky, D.S., and Lewis, D.A. (1998). A subclass of prefrontal gamma-aminobutyric acid axon terminals are selectively altered in schizophrenia. Proc Natl Acad Sci U S A 95, 5341–5346.

Zarrindast, M., Rostami, P., and Sadeghi-Hariri, M. (2001). GABA(A) but not GABA(B) receptor stimulation induces antianxiety profile in rats. Pharmacol Biochem Behav 69, 9–15.

Zhang, B., Seigneur, E., Wei, P., Gokce, O., Morgan, J., and Sudhof, T.C. (2016). Developmental plasticity shapes synaptic phenotypes of autism-associated neuroligin-3 mutations in the calyx of Held. Mol Psychiatry.

Zhang, C., Milunsky, J.M., Newton, S., Ko, J., Zhao, G., Maher, T.A., Tager-Flusberg, H., Bolliger, M.F., Carter, A.S., Boucard, A.A., Powell, C.M., and Sudhof, T.C. (2009). A neuroligin-4 missense mutation associated with autism impairs neuroligin-4 folding and endoplasmic reticulum export. J Neurosci 29, 10843–10854.

Zhou, Y., Kaiser, T., Monteiro, P., Zhang, X., Van der Goes, M.S., Wang, D., Barak, B., Zeng, M., Li, C., Lu, C., Wells, M., Amaya, A., Nguyen, S., Lewis, M., Sanjana, N., Zhou, Y., Zhang, M., Zhang, F., Fu, Z., and Feng, G. (2016). Mice with Shank3 Mutations Associated with ASD and Schizophrenia Display Both Shared and Distinct Defects. Neuron 89, 147–162.

## Supplemental References

1. Liu P, Jenkins NA, Copeland NG. A Highly Efficient Recombineering-Based Method for Generating Conditional Knockout Mutations. Genome Research. 2003;13(3):476–84.

2. Noll S, Hampp G, Bausbacher H, Pellegata N, Kranz H. Site-directed mutagenesis of multi-copy-number plasmids: Red/ET recombination and unique restriction site elimination. BioTechniques. 2009;46(7):527–33.

3. Schneider Gasser EM, Straub CJ, Panzanelli P, Weinmann O, Sassoe-Pognetto M, Fritschy JM. Immunofluorescence in brain sections: simultaneous detection of presynaptic and postsynaptic proteins in identified neurons. Nat Protoc. 2006;1(4):1887–97.

4. Zhao S, Ting JT, Atallah HE, Qiu L, Tan J, Gloss B, et al. Cell type-specific channelrhodopsin-2 transgenic mice for optogenetic dissection of neural circuitry function. Nat Methods. 2011;8(9):745–52.

5. Ting JT, Daigle TL, Chen Q, Feng G. Acute brain slice methods for adult and aging animals: application of targeted patch clamp analysis and optogenetics. Methods Mol Biol. 2014;1183:221–42.

